# Microplastics Disrupt Predator-Induced Plasticity in *Daphnia* across Behavioral, Morphological and Molecular Levels

**DOI:** 10.64898/2026.05.12.724522

**Authors:** Julian Brehm, Marco M Rupprecht, Michael Schwarzer, Luca Liprandi, Anja FRM Ramsperger, Lucia Stuhr, Leon Gasteiger, Lea Bek, Julia Umbach, Jana K Koch, Leon Gröschel, Matthias Schott, Daniel Wagner, Andreas Römpp, Seema Agarwal, Thomas Fröhlich, Christian Laforsch

## Abstract

Microplastics (MP) are widespread in aquatic ecosystems and pose a threat to freshwater biodiversity. While numerous studies examine physiological effects on aquatic organisms, less is known about how MP alter chemically mediated interactions that regulate predator–prey dynamics. Predator-induced defenses in *Daphnia* depend on detecting kairomones and represent an important form of adaptive phenotypic plasticity. Whether MP interfere with these responses, and through which mechanisms, remains unclear. Here, we show that polystyrene MP impair predator-induced defenses across *Daphnia* species by disrupting predator-cue-mediated plasticity at the behavioral, morphological, and molecular levels. In *D. longicephala*, chronic exposure to PS fragments weakened *Notonecta*-induced morphological defenses, whereas additive-containing PS fragments nearly suppressed defense formation and reduced body size. Consistent with these phenotypic effects, proteomic analyses revealed alterations in pathways related to molting and chitin metabolism, linking MP exposure to impaired defense formation. In *D. magna*, PS particles attenuated fish kairomone-induced diel vertical migration, with stronger effects for larger particles, consistent with reduced effective availability or perception of predator cues. Natural limestone particles caused only minor effects, indicating particle-specific rather than general particle-driven responses. Our findings demonstrate that MP can disrupt adaptive predator–prey interactions with potential cascading consequences for freshwater food webs.

## Introduction

Global plastic production is steadily rising, generating persistent waste streams, as most plastics have service lives of less than one year^1,2^. Around 23% of plastic waste is mismanaged, and consequently, plastic debris has become ubiquitous across ecosystems and is projected to increase by up to 30 Mt annually until 2040 if no measures are taken to prevent plastic leakage into the environment^3^. Once released into the environment, plastic debris undergoes continuous fragmentation driven by, e.g., UV irradiation or abrasion, progressively forming microplastic (MP) particles^4,5^. MP particles, most commonly defined by size, for example, as plastic particles of 1 to 1000 μm in size^6^, are now widely documented in freshwater ecosystems worldwide, with particles in the low micrometer range being the most abundant^7,8^.

Once MPs are generated within or enter the environment, they become accessible to a wide variety of organisms. Upon uptake, there is increasing evidence that the biological effects of MP strongly depend on the physicochemical properties of the particles, such as their shape, size, polymer type, and surface properties like their zeta potential or functional groups^9–11^. Further, MP particles found in the environment contain a diverse set of chemical additives, including antioxidants, stabilizers, and plasticizers^12,13^, incorporated during the production process of plastics, to improve material performance and durability^14^. As these additives are not covalently bound to the polymer, but rather physically embedded in the polymer matrix^15^, they can leach into the surrounding environment, a process accelerated by degradation and aging^16,17^. Polymer stabilizers are among the most widely used additives, with concentrations in plastics ranging from 0.05 to 3 wt%, depending on polymer type and additive chemistry^18^. Compared with other additives, phosphite antioxidants such as Irgafos 168 are used in relatively high amounts^18^. Their widespread application has raised concerns about their environmental impact during plastic aging and fragmentation^13^. Consequently, organisms are likely not exposed to a simple, pristine MP particle but rather to a complex particle with varying physicochemical properties, including leachates that may be released upon ingestion or into surrounding media.

Freshwater zooplankton, particularly species of the genus *Daphnia*, are key-stone species in lentic waters. As dominant herbivores in lentic ecosystems, *Daphnia* strongly shape phytoplankton communities and mediate energy transfer to higher trophic levels, thereby shaping food web structure and ecosystem functioning^19^. Hence, they are used as model for assessing effects deriving from environmental pollutants^20^. Modeling indicates that zooplankton are exposed to small MPs throughout the entire stagnation phase in the epilimnion^21^. Consequently, their non-selective filter-feeding behavior leads to high ingestion rates of suspended particles, including MP^22^. Numerous studies reported effects of MP on *Daphnia,* however, most studies primarily focused on direct physiological endpoints like behavior, growth, and reproduction when exposed to MP^11,23–25^. In contrast, comparatively less attention has been given to how MP may alter biotic interactions. In natural ecosystems, fitness depends not only on baseline physiological performance but also on the capacity to respond to ecological challenges such as predation^26^. Predation is a major driver of community dynamics and has led to the evolution of inducible defenses, which are only expressed when organisms detect a reliable chemical cue of predation risk by perceiving so-called kairomones^27^. Accurate cue detection is essential for mounting inducible defenses that balance protection against energetic and developmental costs^28^. Such predator-induced plasticity can be described as a reaction norm, in which defense expression varies predictably along gradients of perceived predation risk^29^. Prey typically respond to kairomones by altering behavior, morphology, and life history^27,30^. One model species that exhibits pronounced inducible morphological defenses in response to kairomones released by predatory backswimmers of the genus *Notonecta*, including the development of large protective crests and increased carapace rigidity^31,32^ is *Daphnia longicephala*. These responses are tightly regulated and energetically costly, with defense magnitude scaling with perceived predation risk^33,34^. On the other hand, *Daphnia magna* is a well-studied model species for inducible behavioral defenses upon exposure to fish kairomones^35,36^, such as diel vertical migration (DVM). DVM, a behavioral pattern where zooplankton move to deeper waters during the day and back to surface waters at night, is primarily controlled by light and activated by chemical cues from fish predators to reduce predation^37^. Because DVM represents a rapid and reversible response to predator detection, it provides a sensitive behavioral proxy for kairomone perception. Together, these systems offer complementary levels at which potential disruptions of predator-induced plasticity can be assessed.

Recent studies suggest that plastic waste and MP particles may weaken the formation and expression of inducible defenses in prey organisms^38–40^, with MP particle size modulating responses^39,40^. However, the available literature lacks mechanistic insights into how MPs interfere with defense induction, whether by reducing fitness through increased energy demands for detoxification processes or by disrupting the detection of chemical cues. Furthermore, these studies have focused on pristine polymers, without considering the role of incorporated additives, and have not used natural control particles to account for the presence of particles *per se*.

To address these knowledge gaps, we investigated whether polystyrene MPs, with and without the antioxidant Irgafos 168, disrupt or weaken chemically mediated predator-prey induced morphological and behavioral defenses. We tested three non-exclusive hypotheses: (i) MP particles reduce kairomone perception by the prey, (ii) MP uptake evokes physiological constraints limiting defense expression, and (iii) additive containing MPs exert additional or synergistic effects via chemical leaching. By including ground mussel shell fragments (limestone) as a natural reference particle, we further tested whether the responses are MP-specific rather than general consequences of particle exposure.

To test these hypotheses, we integrated behavioral assays, phenotypic analyses, proteomic profiling, and chemical characterization within a hierarchical framework. DVM in *D. magna* served as a rapid behavioral proxy for predator cue perception, while inducible morphological defenses in *D. longicephala* quantified changes in defense magnitude under predation risk. Proteomic analyses provided mechanistic insights into physiological alterations that may constrain defense expression. Complementary measurements of Irgafos 168 leaching into the surrounding media at different pH levels were conducted to determine whether additive leaching could account for the observed biological effects.

## Results

### Particle exposure modifies inducible morphological defenses

We measured predator-induced morphological traits in *D. longicephala* exposed to *Notonecta* kairomones (one predator per 600 mL), and to 2 mg/L PS fragments, PS fragments + 0.05% Irgafos 168, and limestone fragments as a natural control, after 14 days of chronic exposure.

Predator cues induced pronounced defensive morphologies, resulting in a strong multivariate effect of predator exposure, i.e., the expression of inducible morphological defenses (Pillai’s trace = 0.93, F₅,₆₂ = 163.8, p < 0.001). Particle treatment also significantly modified predator-induced responses (relative crest height, -width and relative spine length), as indicated by a significant treatment × predator interaction (Pillai’s trace = 0.94, F₁₅,₁₉₂ = 5.88, p < 0.001). The body length, on the other hand, was not affected by predator presence or absence within the treatments.

In the control treatment, predator exposure induced the strongest defensive phenotype. This was mainly reflected by pronounced increases in relative crest height (F₁,₆₆ = 307.5, p < 0.001) and relative crest width (F₁,₆₆ = 499.0, p < 0.001), representing the largest predator-induced changes among all treatments (Figure 1 A, B, & Figure 3). Relative spine length also increased significantly under predator exposure (F₁,₆₆ = 8.98, p = 0.004), although this response was weaker and more variable than the crest-related traits (Figure 2 A & Figure 3).

**Figure 1.**
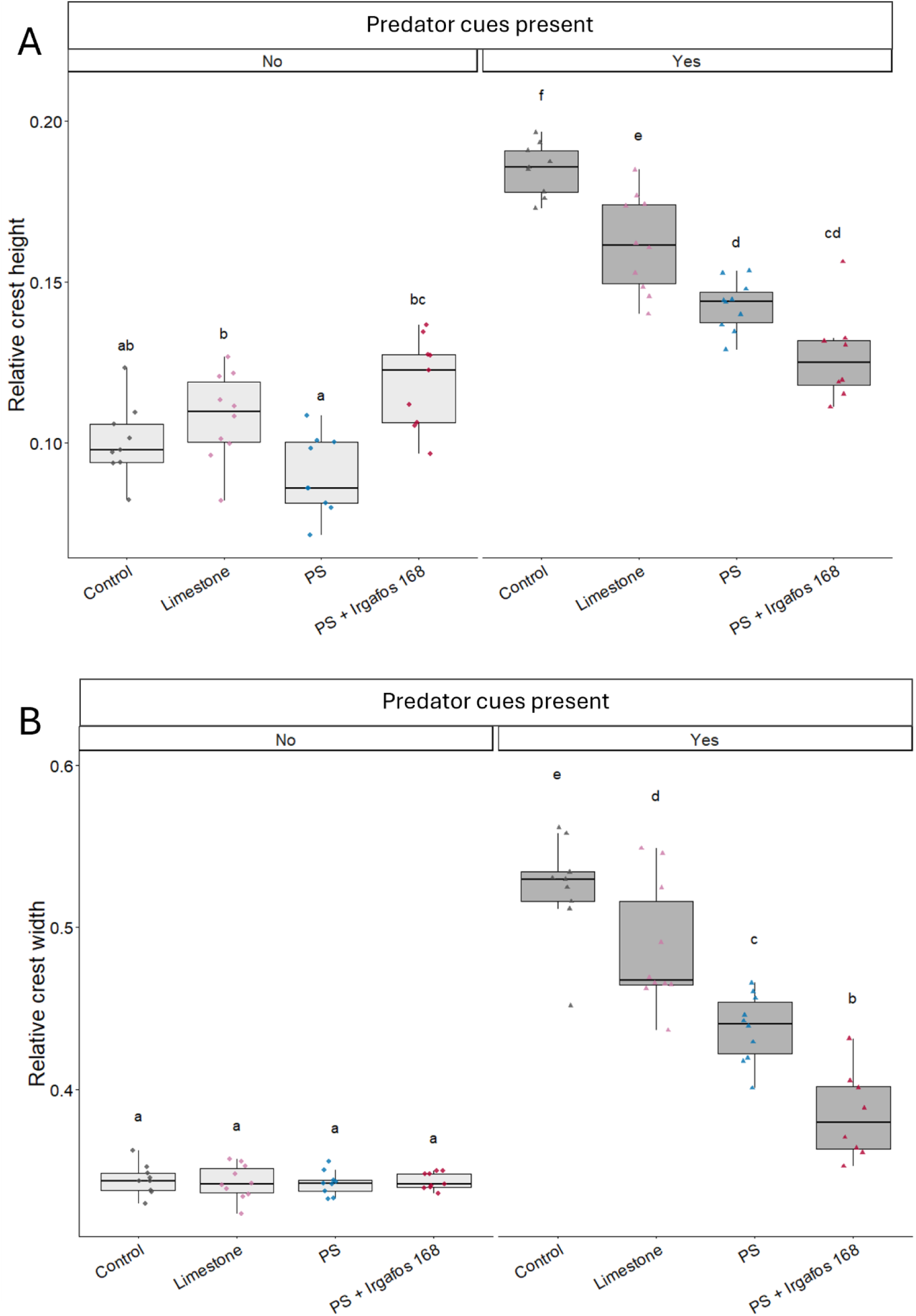
MP particles and associated additives impair predator-induced morphological crest formation in *D. longicephala.* Relative crest height (A), crest width (B) of *D. longicephala* exposed to predator kairomones from *Notonecta* under different particle treatments. Different letters indicate significant differences among treatment × predator groups (Tukey-adjusted post hoc tests, p < 0.05). Boxes represent the interquartile range with the median indicated by the central line; whiskers denote 1.5 × the interquartile range, and points represent individual animals.

**Figure 2.**
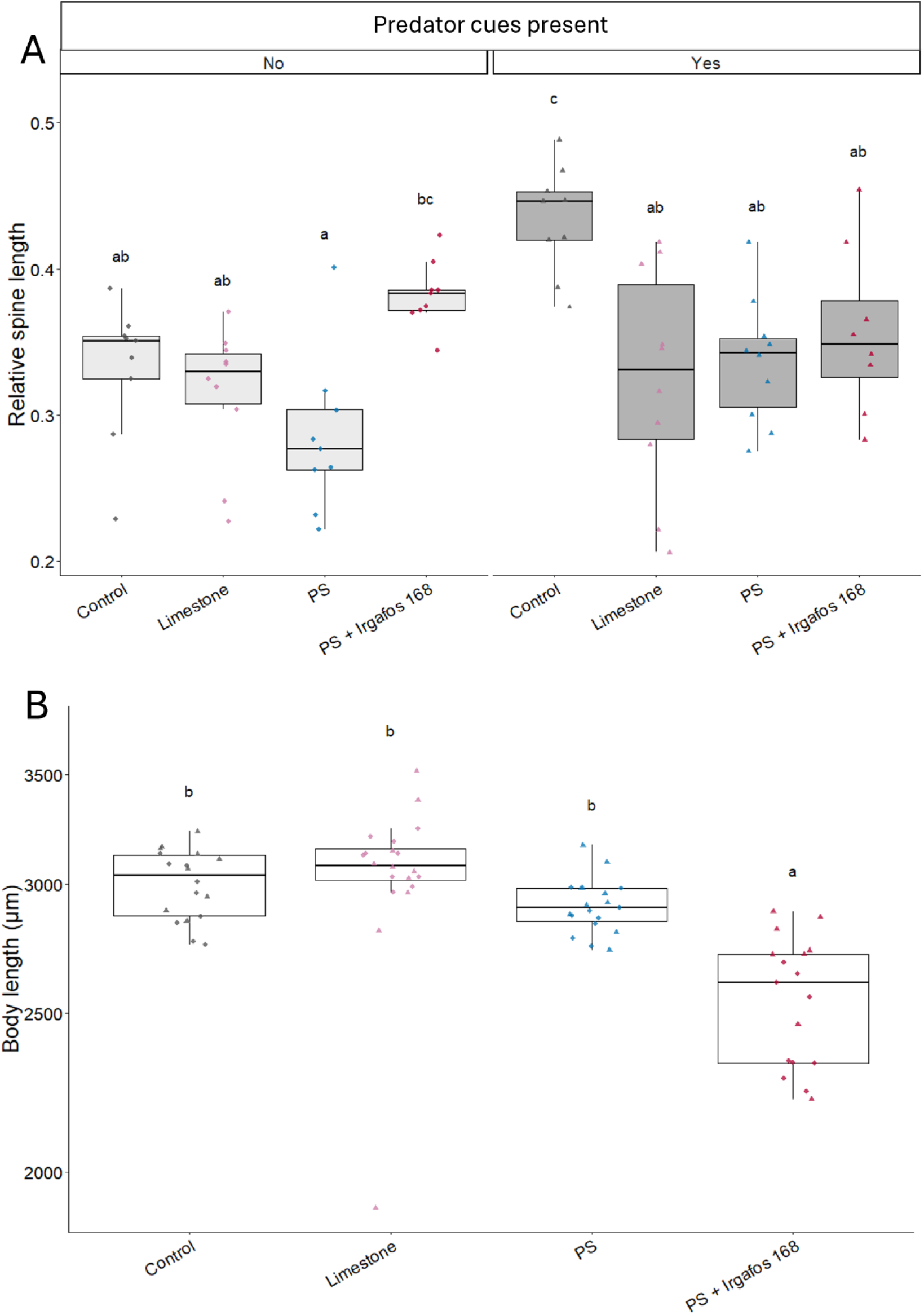
MP particles and associated additives impair relative spine length and body length in *D. longicephala.* Relative spine length (A) of *D. longicephala* exposed to predator kairomones from *Notonecta* under different particle treatments. (B) Body length differed among treatments (squares: predator cues absent; triangles: predator cues present), with individuals exposed to PS + Irgafos 168 exhibiting reduced body size relative to control, limestone, and PS treatments. Different letters indicate significant differences among treatment × predator groups (Tukey-adjusted post hoc tests, p < 0.05). Boxes represent the interquartile range with the median indicated by the central line; whiskers denote 1.5 × the interquartile range, and points represent individual animals.

**Figure 3.**
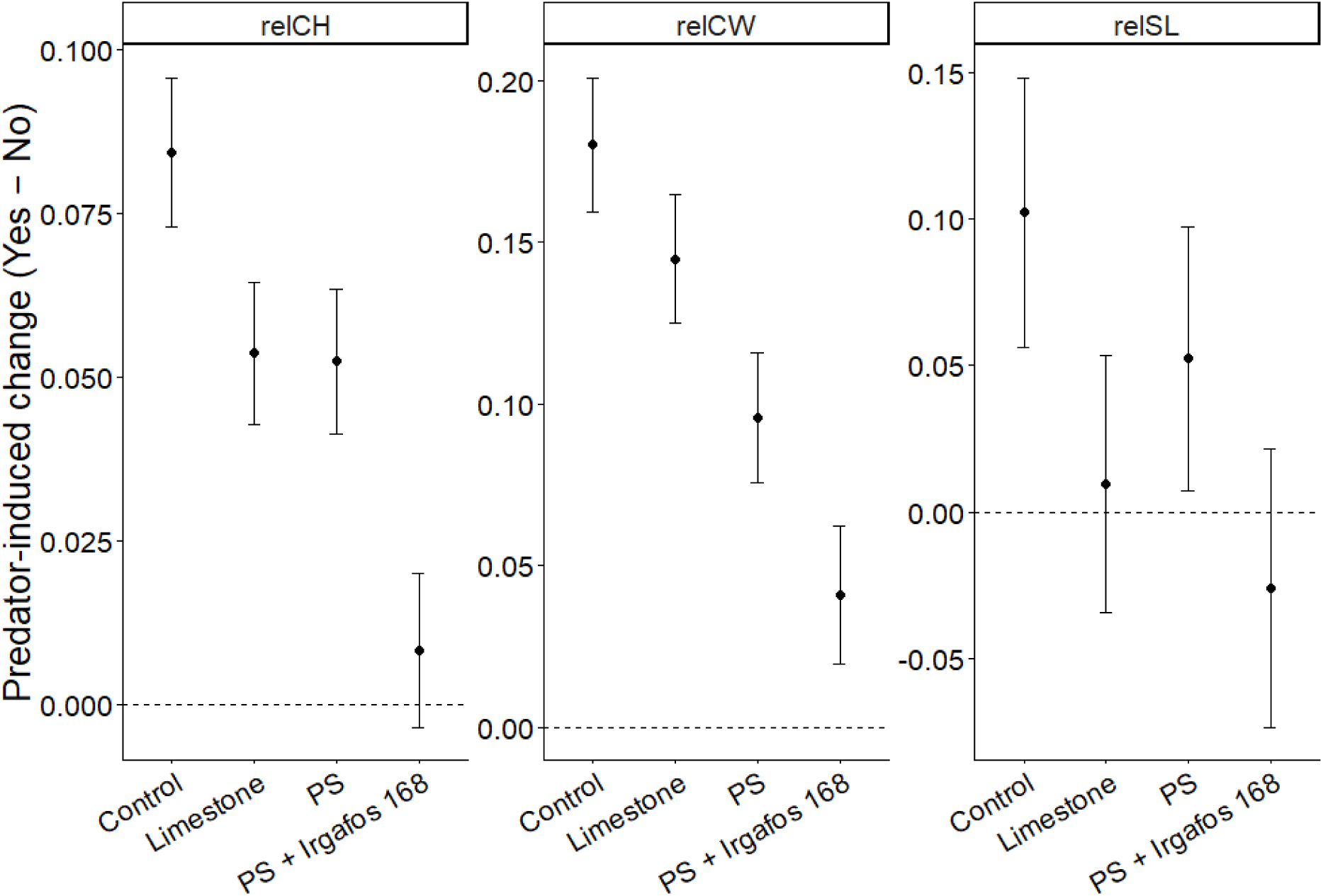
MP particles and associated additives impair predator-induced morphological defenses in *D. longicephala.* Predator-induced change in morphological traits (difference between predator and no-predator conditions) based on estimated marginal means ± 95% confidence intervals (relCH: relative crest height: relCW: relative crest width; relSL: relative spine length).

In the limestone treatment, predator-induced morphological defenses were still clearly expressed, but consistently slightly less pronounced than in the control animals. As shown by the estimated marginal means of predator-induced trait changes (Figure 3), both relative crest height and relative crest width showed smaller predator-induced shifts than in the control treatment, indicating a moderate attenuation of inducible defenses. Relative spine length under predator exposure was significantly lower than in the control treatment (Control vs. Limestone: estimate = 0.109, p < 0.0001), while body length did not differ from control animals (Control vs. Limestone: estimate = -0.011, p = 0.974). Thus, limestone particles caused only a comparatively weak reduction of predator-induced defenses.

In the PS treatment, predator-induced defenses (relative crest height and particularly relative crest width) were more strongly impaired than in the limestone treatment, compared to the control (Figure 1), indicating that PS fragments alone interfered with defense induction. Relative spine length was likewise significantly lower than in predator-exposed control animals (Control vs. PS: estimate = 0.097, p = 0.0004). In contrast to the strong effects on inducible defense traits, body length remained unaffected and did not differ significantly from either control or limestone animals (Control vs. PS: estimate = 0.031, p = 0.633; Limestone vs. PS: estimate = 0.042, p = 0.355), suggesting that PS particles primarily impaired defense expression rather than general growth.

The strongest impairment occurred in the PS + Irgafos 168 treatment. Here, predator-induced morphological defenses (relative crest height and relative crest width) were largely suppressed, resulting in the smallest predator-induced trait changes among all treatments (Figure 1 & Figure 3). Relative spine length also remained significantly lower compared to predator-exposed control animals (Control vs. PS + Irgafos 168: estimate = 0.077, p = 0.012). In addition, PS + Irgafos 168 was the only treatment that significantly reduced body length compared with all other treatments (Control vs. PS + Irgafos 168: estimate = 0.161, p < 0.0001; Limestone vs. PS + Irgafos 168: estimate = 0.172, p < 0.0001; PS vs. PS + Irgafos 168: estimate = 0.130, p < 0.0001; Figure 2 B), indicating that additive-containing particles caused the strongest overall disruption of morphology and inducible defenses.

Together, these results suggest that MP exposure can disrupt predator-induced defense formation in *Daphnia* and indicate potential negative effects on fitness, with MPs containing additives having the most significant impact.

### Proteome analysis

The *D. longicephala* animals of the chronic exposure experiments were further used for proteome analyses, to unravel mechanisms underlying impairment of predator-induced morphological plasticity. Using a DIA-based label-free LC-MS/MS approach, 25723 peptides were identified, which could be assigned to 3361 different proteins (FDR < 1%). After treatment, pronounced proteomic alterations were detected for example for (A) predator cues alone, (B) PS vs. Limestone in presence of predator cues and (C) PS with and without Irgafos 168 in the presence of predator cues (Figure 4 A-C) (see SI Tables 1-3 for more details).

**Figure 4.**
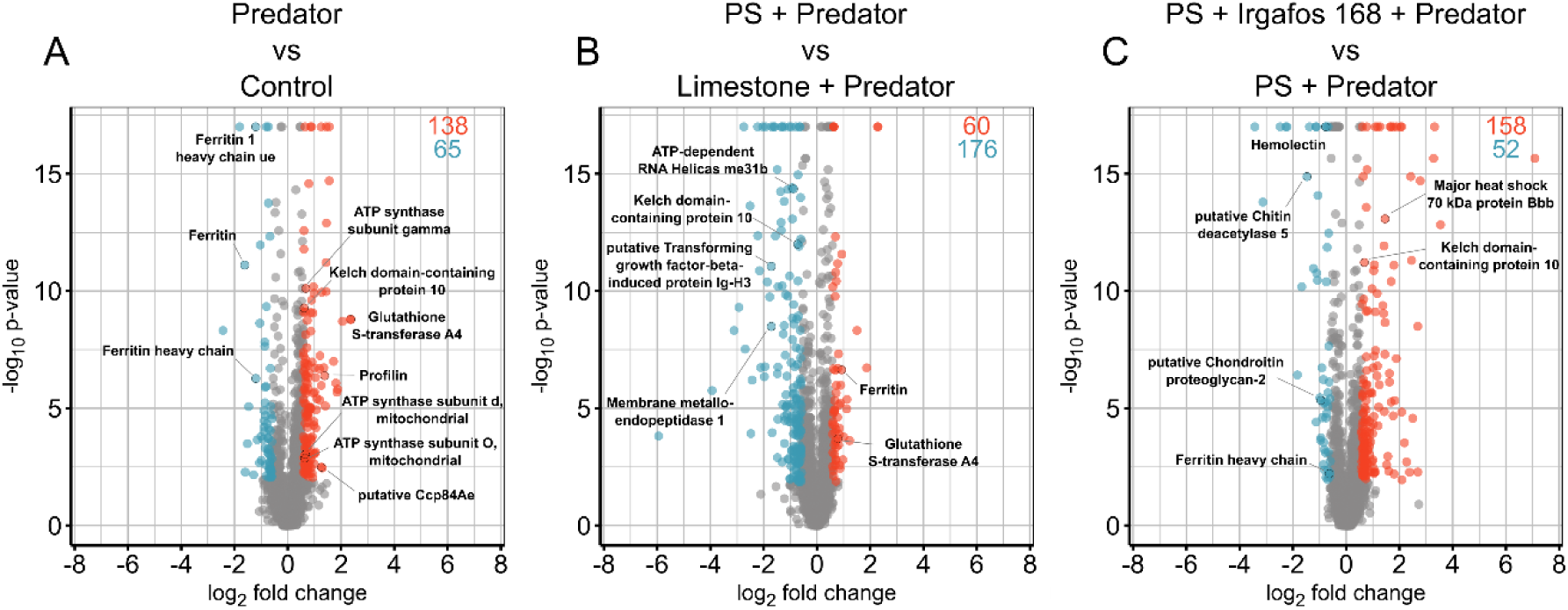
Volcano plots displaying differentially abundant proteins for the respective comparisons. Proteins less abundant are marked in blue, while proteins more abundant are marked in red. Grey proteins are not significantly altered in abundance. (A) Alterations of Predator cues vs. Particle-free control. (B) Alterations of PS + Predator cues vs. Limestone + Predator cues. (C) Alterations of PS + Irgafos 168 + Predator cues vs. PS + Predator cues.

**Table 1:** Overview of the particles used in the respective experiments and their size.

| Particle type | Size [$\mu\text{m}$] | Experiments |
| --- | --- | --- |
| PS spheres | 20 (18.0-22.0) | DVM |
| PS spheres | 45 (40.5-49.5) | DVM |
| PS fragments | $d_{90}$: 16.27 | DVM & Inducible morphological defenses |
| PS fragments + 0.05% Irgafos | $d_{90}$: 17.01 | Inducible morphological defenses & Leaching |
| Limestone fragments | $d_{90}$: 3.01 | DVM & Inducible morphological defenses |

To gain further functional insights, we performed a STRING analysis with the lists of differentially abundant proteins for comparisons (A), (B) and (C). The six most enriched terms are shown in Figure 5.

**Figure 5.**
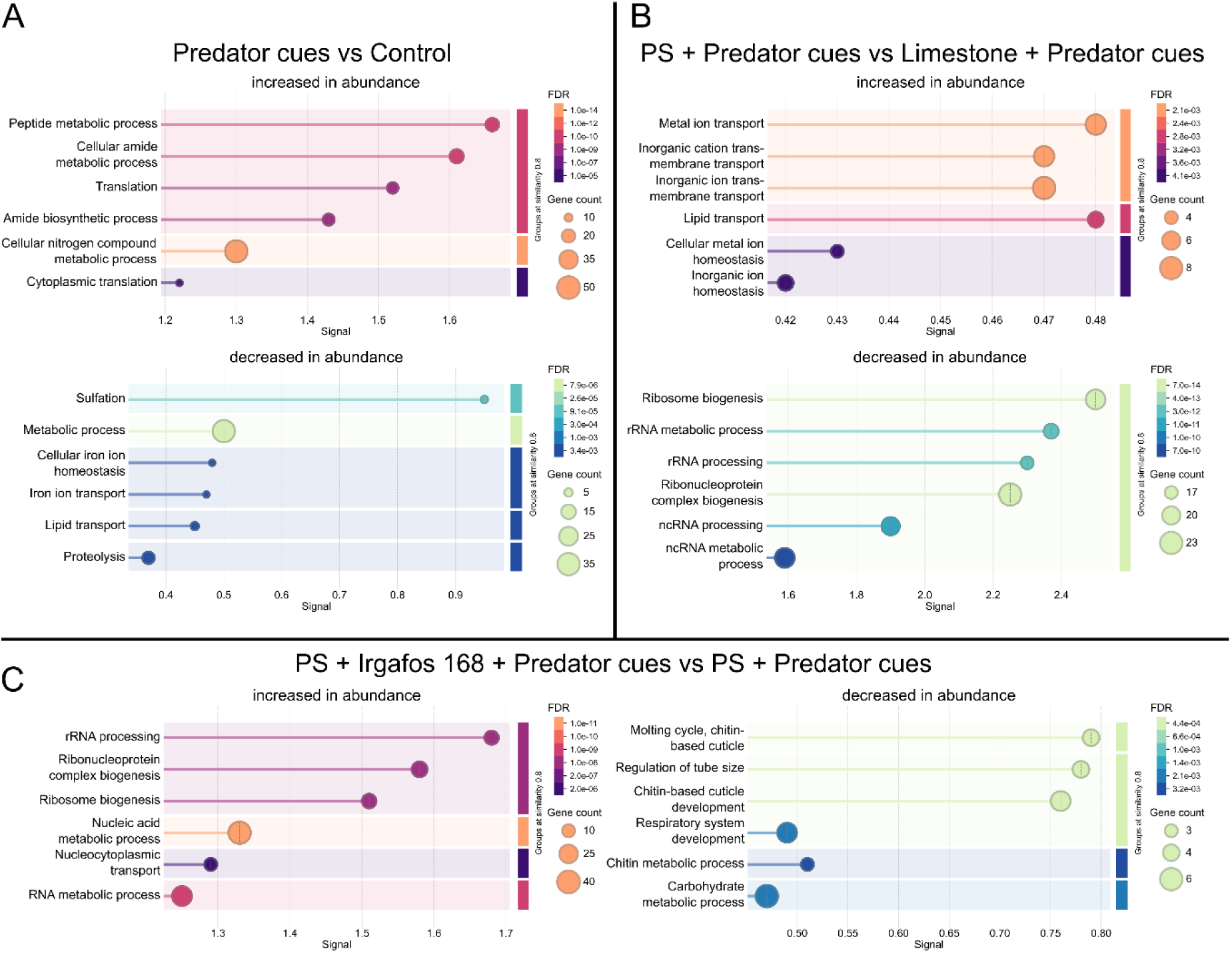
Visualization of the six most significantly enriched Biological Process terms following STRING analysis. Overrepresentation in the set of differentially abundant proteins that were significantly decreased and increased in abundance after exposure to (A) Alterations of Predator cues vs. Particle-free control. (B) Alterations of PS + Predator cues vs. Limestone + Predator cues. (C) Alterations of PS + Irgafos 168 + Predator cues vs. PS + Predator cues. The size of the circles represents the number of differentially abundant proteins within a term, and the significance is encoded as a color gradient.

For predator cues alone (Figure 5A), amongst others, an overrepresentation of proteins increased in abundance, associated with peptide metabolism and translation was found. In contrast, proteins related to sulfation were overrepresented among those that decreased in abundance. Additionally, proteins associated with iron-, lipid transport and proteolysis were decreased in abundance. Regarding the effect of PS in the presence of predator cues compared to limestone (Figure 5B), proteins associated with transport, including those involved in lipid transport and ion homeostasis, were increased in abundance. Notably, PS in the presence of predator cues led to decreased abundances of proteins involved in the regulation of gene expression, particularly ribosome biogenesis and rRNA processing. In contrast, in the presence of predator cues, PS containing Irgafos 168, led to an increased abundance of proteins involved in ribosome biogenesis and rRNA processing when compared to PS without Irgafos 168 (Figure 5C). Conversely, proteins associated with the molting cycle, chitin metabolism, carbohydrate metabolism, and respiratory system development were reduced in abundance in this comparison.

### Leaching of Irgafos 168

To assess whether leaching of the antioxidant additive Irgafos 168 into the media may be a reason for observed effects, PS fragments containing 0.05 wt% Irgafos 168 were incubated in buffer solutions covering a broad pH range (pH 2–10) and analyzed for leachates using GC–MS.

No Irgafos 168 could be detected in extracts obtained from PS particles incubated in aqueous buffers across the tested pH range. Similarly, no Irgafos 168 was detected after Soxhlet extraction under boiling ultrapure water conditions. In all cases, measured concentrations remained below the GC–MS detection limit of 0.5 mg/L for Irgafos 168. These results indicate that either no detectable leaching occurred or that released concentrations remained below the analytical detection threshold. The absence of detectable leaching across all aqueous extraction conditions is consistent with the hydrophobic properties of Irgafos 168.

### Diel Vertical Migration in D. magna

To determine whether suspended particles interfere with the perception or effective availability of predator cues, and thereby alter inducible behavioral defenses, DVM of *D. magna* exposed to fish kairomones from *Leucaspius delineatus* was quantified in a modified plankton organ bioassay under different particle treatments and daylight conditions (Figure 6). Spherical PS particles of two sizes (20 µm and 45 µm in diameter) were used to test whether particle surface area influences the adsorption of kairomones and, accordingly, behavioral responses. Since particle number was kept constant across treatments (1000 P/mL), the larger spheres provided a greater total surface area (6.36 mm^2^/mL) for potential particle-cue interactions compared to the 20 µm ones (1.26 mm^2^/mL). Individuals exposed to 45 µm spherical PS under daylight conditions were positioned significantly higher in the water column compared to control animals (p = 0.014), while other particle treatments showed weaker or non-significant shifts in mean vertical position (Figure 6). Temporal dynamics differed among treatments, with the 20 µm fragment treatment showing a significantly time-dependent downward shift in vertical position (p = 0.042).

**Figure 6.**
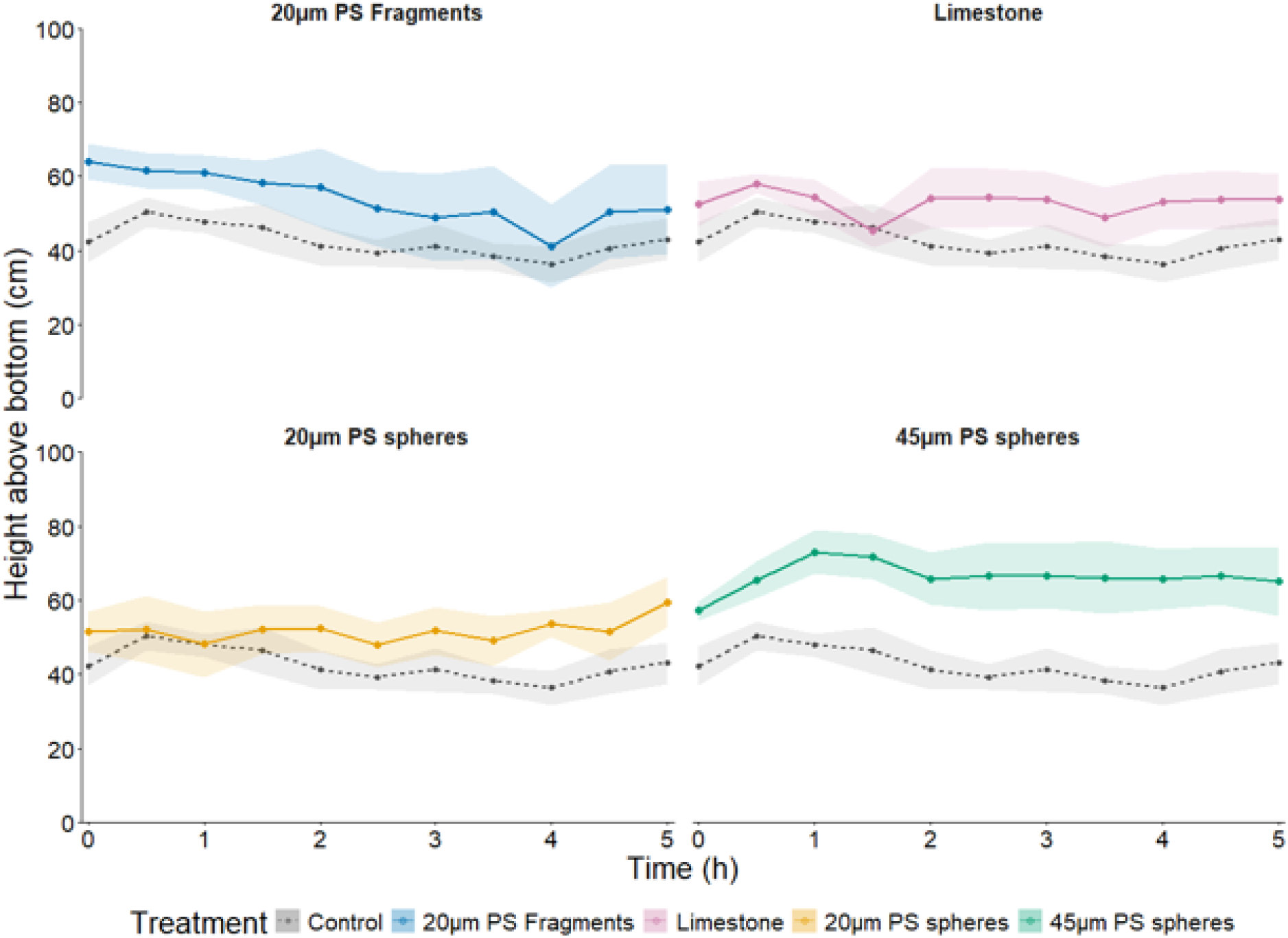
Mean vertical position of *D. magna* over time under predator kairomones (*L. delineatus*), particle treatment, and daylight conditions. Lines represent mean values across replicate tubes (n = 5), and shaded areas indicate mean ± SE. The control response (grey dashed line) is shown in each panel for comparison.

Because predator-induced DVM in *D. magna* represents an avoidance response to visually hunting fish, individuals normally migrate downward upon perception of fish-derived chemical cues, under daylight conditions. The observed shift towards higher positions in the presence of larger particles therefore suggests a reduced perceived predation risk. Given that particle number was identical across treatments, the stronger behavioral effect observed for 45 µm particles is consistent with an influence of particle surface area on the effective perception of predator cues.

## Discussion

We showed, using a systematic approach, that MP particles interfere with predator-induced defenses in *Daphnia* across multiple biological levels. In *D. longicephala*, morphological defense traits, particularly crest height and width, were strongly reduced under MP exposure, especially in the presence of additive-containing MP particles. These phenotypic responses were accompanied by substantial proteomic alterations, giving insight into specific molecular pathways underlying these effects, such as chitin metabolism involved in molting. In addition, MPs impaired DVM behavior in *D. magna*, likely through adsorption of kairomones on the particle surface, leading to reduced effective perception of predator cues.

The progressive reduction of inducible defenses from limestone to pristine PS and additive-containing PS suggests that MPs interfere with predator-induced phenotypic plasticity in *Daphnia*. This disruption may arise from reduced effective availability of predator cues in the presence of suspended particles, impaired chemosensory perception, or downstream physiological constraints affecting defense formation. Because crest formation represents a highly regulated and energetically costly response to *Notonecta* kairomones^31^, the near-complete suppression observed for additive-containing particles points to a substantial disruption of adaptive phenotypic plasticity. Trait-specific sensitivity, which leads to different dose-response patterns of defensive traits^41^ in *Daphnia*, further supports the interpretation that MPs specifically interfere with adaptive defense formation: crest traits, closely associated with predator defense effectiveness against *Notonecta*^42^, were strongly suppressed, while spine length exhibited weaker and more variable responses (Fig. 2 A).

Importantly, these pronounced effects were observed at a particle concentration of 2 mg/L, which exceeds most currently reported background levels in freshwater systems but remains within the range of concentrations considered environmentally relevant for studying MP effects in freshwater^43^. While environmental measurements are often reported in particle numbers and are therefore difficult to directly compare to mass-based concentrations, available literature suggests that µg L⁻¹ to low mg L⁻¹ levels are reached in highly impacted marine and freshwater habitats^44,45^. Consequently, the exposure level used here should be interpreted as representative of elevated or worst-case scenarios, which may become increasingly relevant under continued plastic emissions and environmental accumulation^46^.

The progressive impairment of inducible defenses was accompanied by distinct treatment-specific proteomic signatures, providing mechanistic insight into the underlying physiological alterations. Notably, MPs containing 0.05% Irgafos 168 induced a distinct proteomic signature relative to PS exposure alone under identical predator conditions, which closely corresponds with the near-complete suppression of inducible defenses. This progressive pattern, ranging from natural particles to pristine PS and finally additive-containing PS, closely mirrored the pattern of the observed morphological effects. In the presence of Irgafos 168, proteins involved in ribosome biogenesis and rRNA processing were increased in abundance, indicating a shift in the regulation of translational processes. In parallel, proteins such as major heat shock 70 kDa protein Bbb (KZS19529.1) were elevated, suggesting activation of the cellular stress response^47^. However, this was accompanied by an overrepresentation of abundance-reduced proteins related to the molting cycle, chitin metabolism, and carbohydrate metabolism (Fig. 5C). Since the expression of morphological defenses in *Daphnia* relies on chitin-based cuticular structures and inducible morphological defenses are formed from molt to molt^48^, the downregulation of these pathways provides a mechanistic link to the observed reduction in crest height and width. Consistent with these physiological alterations, individuals exposed to additive-containing PS fragments also exhibited reduced body size, a trait closely linked to fecundity and overall fitness^49,50^. Together, these findings suggest that additive-containing particles impose substantial physiological constraints extending beyond defense formation alone. Although no detectable leaching of Irgafos 168 into the aqueous phase was observed under the tested conditions, concentrations may still have remained below analytical detection limits. Therefore, the observed effects may either result from very low-level additive release, direct particle-associated interactions, or localized interactions at the particle-organism interface, i.e., leaching in the gut.

While additive-containing particles caused the strongest molecular disruption, PS particles without additives already altered proteomic responses relative to natural particles under identical predator conditions. This shift was characterized by an increased abundance of proteins associated with ion transport, lipid transport, and cellular homeostasis, alongside elevated ferritin (decreased in predator cues vs. control) (KZS12270.1) and Glutathione S-transferase (GST) (KZS14637.1) levels, suggesting adjustments in membrane-associated processes, ion-, and redox balance. Similar changes in metabolic and stress-related pathways have been reported in *Daphnia* exposed to environmental stressors such as cadmium^51^, organically contaminated stream water^52^, and temperature changes^53^. At the same time, proteins associated with biosynthetic and structural capacity, including ATP-related proteins (KZS10902.1, KZS20545.1, KZS20408.1), Kelch domain-containing protein 10 (KZS21807.1), and transforming growth factor beta-induced protein Ig-H3^54^ (KZS21360.1), were reduced in abundance. Consistent with this, processes related to ribosome biogenesis and rRNA processing were significantly reduced, indicating a suppression of translational capacity. As ribosome biogenesis represents a major energetic investment and is closely linked to growth and developmental plasticity^55^, its downregulation suggests a shift in resource allocation away from biosynthesis toward maintenance-related processes^56^. Importantly, these changes occur in direct comparison to individuals exposed to natural particles under predator conditions, indicating that PS modifies the physiological response to predator cues rather than simply inducing a general stress response. This interpretation is consistent with the observed attenuation of inducible defenses at the morphological level. Crest formation in *D. longicephala* requires rapid structural remodeling^57^, and thus depends on sufficient biosynthetic capacity. The observed reduction in translational machinery, therefore, provides a plausible mechanistic explanation for the impaired expression of these traits.

To place these treatment-specific changes into the context of inducible defense formation, we next examined the proteomic response to predator cues alone. Exposure to predator cues alone led to increased abundance of proteins related to peptide metabolism and translation, alongside enrichment of processes related to amide biosynthesis and cytoplasmic translation, indicating an enhanced translational/anabolic capacity. This response was accompanied by increased abundance of ATP synthases (KZS18123.1, KZS13149.1, KZS11497.1), consistent with possible elevated energetic demands for inducible defenses^58^, and proteins such as profilin (KZS10851.1), associated with cytoskeletal dynamics in all eukaryotes^59^, and putative Ccp84Ae (KZS12048.1), a structural constituent of cuticle, which may support structural remodeling during defense formation. In addition, several GSTs (KZS14637.1, KZS11037.1, KZS19194.1, KZS15273.1) were elevated, which in *Daphnia* are usually increased in response to biotic^60^ and abiotic^61^ stressors, reflecting adjustments in redox balance and cellular homeostasis commonly associated with general oxidative stress response^62^. On the other hand, the abundance of ferritin (KZS12270.1) and its heavy chain subunits (KZS17638.1, KZS12271.1) were reduced, together with processes related to iron ion homeostasis and transport. This decrease may reflect a reorganization of iron metabolism, as ferritin can be degraded under increased cellular demand to release iron for essential metabolic processes^63^. The decrease in the abundance of multiple proteins, which are associated with proteolysis, further suggests a shift in protein homeostasis. However, direct evidence for altered protein degradation during inducible defense formation in *Daphnia* remains limited. Similar reductions in proteolytic components have been reported under abiotic stress conditions such as salinity in *Daphnia pulex*^64^, indicating that proteostasis may be reorganized under changing environmental conditions. Taken together, the increase in translational and biosynthetic processes, elevated energy production, and the reorganization of iron metabolism and protein turnover reflect a metabolic adjustment that is consistent with molecular signatures previously associated with inducible defense formation in *Daphnia*^58,65^. Hence, the observed changes likely support the rapid structural remodeling required for morphological defenses, providing a mechanistic link between molecular responses and phenotypic plasticity.

Since inducible defenses in *Daphnia* rely on reliable predator-cue perception, we next examined whether suspended MPs also interfere with predator-induced behavioral responses. We quantified the DVM induction in *D. magna* under predator-cue conditions and daylight. Our results indicate that suspended MP particles can attenuate predator-induced behavioral responses, as individuals exposed to larger (45 µm) spherical PS particles exhibited a reduced downward migration compared to controls, limestone, and other PS particles. As downward migration during the day is a well-established response to fish cues^36^, this shift indicates a reduced perceived predation risk. Our findings of reduced migration are consistent with previous work demonstrating that DVM in *Daphnia* is tightly regulated by chemical predator cues and highly sensitive to changes in cue concentration and reliability^66–68^. This suggests that suspended MPs may reduce the effective availability of kairomones, potentially through adsorption to particle surfaces^38^, thereby weakening predator-cue signaling and altering predator-induced behavioral responses. Taken together, our data indicate that suspended MP particles can reduce the availability and perception of predator kairomones in a surface-dependent manner, potentially through adsorption to particle surfaces and interference with chemosensory signaling. This may lead to a mismatch between actual predation risk and behavior^69^. An effect also observed for environmentally relevant concentrations of copper and nickel in *Daphnia*^69^. Taking into account that DVM acts as a biological nutrient and carbon pump^70^ and that *Daphnia* act as the dominant link between phytoplankton and fish in lake food webs^71^, a disrupted DVM may have implications for trophic interactions.

In summary, the detected proteomic alterations align with the observed impairment of inducible morphological defenses. In particular, the altered abundance of proteins associated with molting, and chitin metabolism suggests that MPs constrain physiological processes required for defense formation. These molecular changes are consistent with the reduced crest development observed under PS exposure, especially in the presence of additive-containing particles. In parallel, the altered DVM behavior in *D. magna* indicates that suspended MPs may reduce the effective availability or perception of predator cues, potentially through adsorption of kairomones to particle surfaces. Together, these findings suggest that MPs interfere with predator-induced plasticity both by affecting predator-cue signaling and by constraining the physiological processes underlying defensive trait formation. Since it is essential for *Daphnia* to adapt its defenses to current predation risk, maladaptation of defensive traits when simultaneously exposed to MP may have knock-on effects on population dynamics across multiple trophic levels, as *Daphnia* is a key species in lentic ecosystems.

## Methods

### Particle preparation and characterization

PS MP spheres were purchased from Polysciences Europe GmbH (Hirschberg an der Bergstrasse, Germany) (Polybead Polystyrene Microsphere 45.0 µm (Article Nr. 07314-5) and Polybead Polystyrene Microsphere 20.0 µm (Article Nr. 18329-5)). PS MP fragments were produced in-house from pellets of the original material, Styrolution PS 158N/L (Ineos Styrolution Group GmbH, Frankfurt am Main, Germany), using a centrifugal mill (ZM 200, Retsch GmbH, Haan, Germany) and were subsequently air jet sieved (e200 LS, Hosokawa Alpine, Augsburg, Germany) to gain fragments smaller than 20 µm. To produce the PS MP fragments containing 0.05% Irgafos 168, a hydrolytically stable phosphite processing stabilizer and secondary antioxidant, the Styrolution PS 158N/L pellets were pre-milled into fragments <1500 μm and dried under vacuum at 40 °C. Then the fragments were mixed in a LM 2.5 powder mixer (MTI Mischtechnik, Detmold, Germany) for 10 min at 500 rpm with 0.05% Irgafos 168 (BASF SE, Ludwigshafen am Rhein, Germany). Subsequently, the obtained mix was extruded with a P11 twin-screw extruder (Thermo Fisher Scientific, Schwerte, Germany) through a 2.2 mm nozzle at 220 °C and 110 rpm. Lastly, the extruded mix was milled under the same conditions as the pristine PS MP fragments to obtain fragments similar in size (< 20 µm). As natural control particles, limestone fragments were used. These were obtained from cautiously cleaned shells from the freshwater mussel *Dreissena bugensis*. The shells were collected in 70% ethanol, then scrubbed with wooden brushes with natural bristles and rinsed several times with ddH2O. After drying, the milling and sieving process used for the PS MP fragments was applied to obtain fragments <20 µm.

Laser diffraction (LD) and dynamic image analyses (DIA) were performed (FLOWSYNC, Microtrac MRB, Montgomeryville and York, USA) for all milled fragments to obtain d_(90)_ values (Table 1).

To obtain further information on the morphology of the particles, scanning electron microscopy (SEM) was conducted (see Figure S1). To visualize the pristine spherical MP from the DVM experiments, the stock dispersion was diluted 1:100 in ultrapure water (UltraPure^TM^, Distilled Water, DNase/RNase Free, Invitrogen) and centrifuged for 20 min at 2000 x g. The supernatant was removed by pipetting, and the pellet was redispersed in 20 µL ultrapure water. The MP dispersions were pipetted onto a glass coverslip (diameter: 12 mm, #1, MENZEL GLAESER, Braunschweig, Germany), placed on carbon conductive tabs (Ø 12 mm Plano GmbH, Wetzlar, Germany), fixed to aluminum stubs (Ø 12 mm, Plano GmbH, Wetzlar, Germany), and left to dry in a desiccator.

The particles used in the experiment to evaluate inducible morphological defenses were prepared for SEM in their pristine form. Therefore, 10 µL of ultrapure water was pipetted onto the glass coverslips mounted on the aluminum stubs as described above, and a spatulaful of the MP powder and the respective natural control particles was mixed with the ultrapure water and left to dry in a desiccator. Before imaging was conducted using a JEOL JSM-IT500 Scanning Electron Microscope (SEM, JEOL Ltd., Japan) each sample was sputter coated with 4 nm Platinum (Leica EM ACE600, Leica Microsystems GmbH, Germany).

### Irgafos 168 GC-MS measurements in water

To investigate the potential release of Irgafos 168 from PS particles, 1 g of PS fragments containing 0.05 wt% Irgafos 168 was incubated in 10 mL buffer solutions at pH 2, 6, 8, and 10 for one week at room temperature under continuous stirring. Following incubation, particles were removed by filtration, and the aqueous phase was extracted with 5 mL dichloromethane (DCM). To additionally test additive release under harsher extraction conditions, 1 g of the same PS particles was extracted in 100 mL boiling ultrapure water (Milli-Q) for 24 h using a Soxhlet apparatus. The aqueous phase was subsequently extracted with 50 mL DCM. Organic extracts were analyzed by gas chromatography–mass spectrometry (GC–MS) using an Agilent 7890B GC system coupled to a quadrupole mass spectrometer detector (Agilent Technologies GmbH, Oberhaching, Germany). Separation was performed on a Zebron ZB-5ms column (5% phenyl methyl siloxane, 95% dimethylpolysiloxane; 30 m × 250 μm × 0.25 μm) using helium as carrier gas at a flow rate of 1 mL/min. Samples were introduced via split injection (1:30) at an inlet temperature of 280 °C. The detection limit for Irgafos 168, determined using analytical standards of known concentration, was 0.5 mg/L.

### Daphnia culture conditions

For the DVM experiments, the laboratory-cultured *D. magna* clone K_3_4J, which originates from a former fishpond near Munich, Germany, was used. Prior to the experiments, 20 female animals were reared in 1.5 L jars containing 1 L of M4 medium^72^. For the experiments on inducible morphological defenses, the laboratory-cultured *D. longicephala* clone Sonja was used. This clone originates from Lara Pond, an urban pond on the periphery of Melbourne, Australia. Prior to the experiments, 20 female animals were reared in 1.5 L jars containing 0.5 L M4 medium and 0.5 L ADaM (“Aachener Daphnien Medium”^73^, with a slightly modified SeO2 concentration (70 mg/L)). In both experiments, the third brood was used, and the first- and second-brood neonates were removed within 24 hours after release to avoid crowding effects. All *Daphnia* were cultured in a climate chamber at 20 °C and maintained a 16:8 h day/night rhythm, with a 30-minute twilight period at dawn and dusk. Daphnids were fed *ad libitum* with the unicellular algae *Acutodesmus obliquus* three times a week.

### Inducible morphological defenses experiment

*D. longicephala* were exposed to eight treatment groups, which were categorized into predator-free and predator-cue treatments. Predator-cue treatments consisted of 600 mL medium conditioned by one *N. glauca* (sampled in small ponds at the University of Bayreuth). All media consisted of equal parts M4 medium and ADaM. Next to particle-free controls, limestone fragments, PS fragments, and PS fragments + 0.05 % Irgafos 168 were used (Table 1). For the six particle treatments, 20 mg particles were added to 10 L medium, corresponding to a final particle concentration of 2 mg/L.

Prior to the experiment, one 12 L aquarium per treatment was prepared, each containing 10 L (5 L ADaM + 5 L M4) prefiltered medium (retention 2-3µm, ROTILABO® Typ 115A, Ø: 90 mm, Carl Roth GmbH + Co. KG, Karlsruhe, Germany). After the addition of the particles, the aquaria were stirred daily with a glass stick, and the particles were allowed to disperse in the medium for 5 days in a climate chamber at 20 ± 0.5 °C.

One day before the start of the experiment, and on each of the following days, 600 mL of the medium was transferred from each aquarium into separate 1.5 L glass beakers. To foster the release of kairomones in the kairomone treatments, one *N. glauca* (body length: 2 ± 0.3 cm) and ten adult *D. longicephala* (serving as prey) were added into the respective beakers. In the four predator-free treatments, only five *D. longicephala* individuals were added to each beaker to maintain comparable medium conditions. After 24 hours, the *Notonecta*, the alive daphnids, and the empty carapaces were removed from the media, and 50 ml portions were distributed into the final experimental beakers. Then, the pooled neonates born within 24h were randomly distributed across the experimental beakers. Media was renewed daily following the described procedure, and *Daphnia* were fed with *A. obliquus* at a concentration of 2 mg C/L throughout the experiment. After 14 days of exposure, the animals’ morphological traits were analyzed under a stereomicroscope equipped with an integrated digital camera (Olympus D26) using CellSens software (v 1.11, Olympus) (Figure 1 & Figure 2). Subsequently, animals were sampled for proteomic analyses.

### DVM experiments

DVM behavior of *D. magna* was quantified using a modified plankton organ bioassay, described previously^74^ in a climate chamber at 20 ± 0.5 °C. This setup consisted of 30 vertically oriented glass tubes (length: 1 m, diameter: 2 cm, closed at the bottom), separated by black partitions to minimize lateral light scattering. Tubes were illuminated only from above with a cold-white LED strip (6000 K) to simulate daylight conditions.

To generate medium containing predator cues, two aquaria were prepared three days before the experiment, each containing 3.5 L of M4 medium and four adult Leucaspius delineates (6±1 cm) that had been pre-fed *D. magna*. After two days, one additional fish and 0.5 L of M4 were added to each aquarium. On the third day, the kairomone-containing media were pooled and filtered to remove any particulates (Whatman™ Grade 3 filter paper, Cytiva, Marlborough, USA) before use in the experiment.

MP and Limestone fragments were suspended in the kairomone-containing media for 1 h before the tubes were filled (1,000 particles/mL). Then, glass tubes were filled with the kairomone-containing media. Five replicate tubes were used per treatment and arranged randomly within the setup. Subsequently, five randomly selected adult *D. magna* were introduced into each tube and fed with 1 mg C/L with *A. obliquus*. The experiment lasted 5 h. During this period, the vertical distribution of individuals was recorded photographically every 30 min at a depth resolution of 5 cm. Mean residential depth was determined from the images.

### Statistics

For the DVM experiment, mean residential depth was analyzed using linear mixed-effects models implemented in the R package lme4^75^. Treatment and time were included as fixed effects, with time modeled as a second-order polynomial to allow for potential non-linear migration dynamics. Tube identity was included as a random intercept to account for repeated measurements of the same experimental units. Observations were weighted by the number of individuals per tube. Significance of fixed effects was assessed using Type III ANOVA with Satterthwaite approximation of degrees of freedom (lmerTest^76^). Treatment effects on vertical position were evaluated using estimated marginal means (emmeans^77^) with Tukey-adjusted pairwise comparisons. Model assumptions were assessed using simulated residual diagnostics implemented in the DHARMa package^78^.

Morphological traits were analyzed using a combination of multivariate and univariate approaches. To assess whether particle exposure and predator cues affected overall morphology, a multivariate analysis of variance (MANOVA) was conducted using the variables body length, relative spine length, relative body width, relative crest height, and relative crest width. Relative traits were calculated by dividing each morphological measurement by body length to account for differences in body size. Significance of multivariate effects was evaluated using Pillai’s trace.

To analyze individual morphological traits, linear models were fitted with treatment, predator cue presence, and their interaction as fixed factors. Model assumptions were assessed using simulated residual diagnostics implemented in the DHARMa package^78^, including tests for uniformity and dispersion as well as visual inspection of residual plots. Residual distributions indicated acceptable model fits for all traits.

Body length exhibited heteroscedastic residuals and was therefore analyzed separately using generalized least squares models implemented in the nlme package^80^. Body length was log-transformed prior to analysis to improve normality, and heteroscedasticity was accounted for by applying a power variance structure. Treatment, predator cues, and their interaction were included as fixed effects.

Post hoc comparisons among treatments were performed using estimated marginal means calculated with the emmeans package^77^. Pairwise comparisons were adjusted using Tukey’s method to control for multiple testing. For traits showing significant treatment × predator interactions, treatment contrasts were evaluated separately within predator cue conditions.

Graphical representations were generated using ggplot2^79^, with boxplots showing trait distributions and overlaid individual observations.

### Proteomics

*D. longicephala* (at least n = 3 per treatment) were washed three times in phosphate buffer (25 mM, pH 7.4), transferred individually into 1.5 mL vials, snap-frozen in liquid nitrogen and stored at -80°C. Daphnids were lysed in 8M Urea, 50 mM ammonium hydrogen carbonate supplemented with protease inhibitors c0mplete ULTRA Tablets, Mini, EASYpack (Roche, Germany), followed by sonication using a Sonopuls HD3200 cup resonator (BANDELIN electronic GmbH & Co. KG, Berlin, Germany) (10 seconds pulse, 20 seconds rest, 12 cycles) and further homogenized using QIAshredder (Qiagen, Hilden, Germany) devices (20817 x g, 1 min). Protein concentrations were determined using the Pierce 660 nm Protein Assay (Thermo Fisher Scientific, Rockford, IL, USA). Proteins were reduced in 4 mM dithiothreitol (DTT) and 1.8 mM tris(2-carboxyethyl)phosphine at 56°C for 30 minutes and then alkylated in 8 mM iodoacetamide at room temperature (RT) for 30 min in the dark. The reaction was quenched with 10 mM DTT at RT for 15 min in the dark. Subsequently, all samples were sequentially digested, first with Lys-C (enzyme-to-protein ratio 1:100, 4 h at 37°C, FUJIFILM Wako Chemicals Europe GmbH, Neuss, Germany), followed by trypsin (enzyme-to-protein ratio 1:50, 18 h at 37°C, Promega, Madison, WI, USA) after adjusting the concentration to 1 M Urea. The digestion was stopped by adding formic acid to a final concentration of 1%. Peptides were dried in a vacuum concentrator and subsequently reconstituted in 0.1% FA. Peptides were desalted using StageTips C18 (Thermo Fisher Scientific, Dreieich, Germany).

LC-MS/MS analysis was performed on a nanoElute 2-LC system connected to a timsTOF HT mass spectrometer (both Bruker Daltonics Inc., Fremont, CA, USA), operated in the data-independent acquisition (DIA) mode. 550 ng of peptides were first loaded onto a trap column (PepMap Neo trap column (300 µm× 5 mm, 5 µm particles, C18, Thermo Fisher Scientific, Waltham, MA, USA)) and separated on a PepSep Ultra C18 25 cm x 75 µm, 1.5 µm (Bruker Daltonics Inc., Fremont, CA, USA), at 250 nL/min with a 25 min gradient of 2-25% of solvent B followed by a 12 min increase to 37%. Solvent A consisted of 0.1% formic acid in water and solvent B of 0.1% formic acid in acetonitrile. The mass spectrometer was run in the dia-PASEF mode using 21 x 25 m/z wide windows and ion mobility ramps between 0.85 and 1.27 1/k0. The sample order was randomized, and carryover was minimized by running blanks between samples. Protein identification and peptide quantification was carried with DIA-NN (v2.2.0), using a fasta file, based on the NCBI *Daphnia magna* genome assembly daphmag2.4 (https://www.ncbi.nlm.nih.gov/datasets/gene/GCA_001632505.1/) (based on Don Gilbert *Daphnia magna* Annotation 1 (November 17, 2022)), alongside the built-in contaminants fasta file. The FDR confidence was set to < 1%, and proteins were filtered for a minimum of two unique peptides. To identify differentially abundant proteins, MS-EmpiRe^81^ (https://github.com/zimmerlab/MS-EmpiRe) was used.

To correct for multiple testing, the false discovery rate was set to be < 0.05 using the Benjamini-Hochberg procedure. All statistical evaluation and bioinformatics, as well as plotting (volcano- and bubbleplots), were performed using R Version 4.5.1^82^. For overrepresentation analysis, STRING (V.12)^83^ and the “biological process” GO database was used. The mass spectrometry proteomics data have been deposited to the ProteomeXchange Consortium (http://proteomecentral.proteomexchange.org) via the PRIDE partner repository^84^ with the project accession PXD076506.

## Ethics approval and consent to participate

Not applicable.

## Consent for publication

Not applicable.

## Availability of data and materials

All data generated or analyzed during this study are included in this published article and its supplementary information files. The mass spectrometry proteomics data have been deposited to the ProteomeXchange Consortium (http://proteomecentral.proteomexchange.org) via the PRIDE partner repository^84^ with the project accession PXD076506.

## Competing interests

The authors declare that they have no competing interests.

## Funding

This study was funded by the Deutsche Forschungsgemeinschaft (DFG; German Research Foundation) - Project Number 391977956 –SFB 1357. The SEM was funded by the DFG (DFG GZ: INST 91/427–1 FUGG). The instrumentation for proteomics was partly funded by the Deutsche Forschungsgemeinschaft (DFG, German Research Foundation; Project Number 534046536).

## Acknowledgments

We thank the technical assistants in the Chair of Animal Ecology I at the University of Bayreuth for their support with animal maintenance and algal culture.

## Notes

http://proteomecentral.proteomexchange.org

### Competing Interest Statement

The authors have declared no competing interest.

