## Supplemental Figure S1 for "Microplastics Disrupt Predator-Induced Plasticity in *Daphnia* across Behavioral, Morphological and Molecular Levels"

* Corresponding author

*Scanning electron micrographs of the used particles*

Scanning electron microscopy analysis of MP and natural control particles showed that the PS spheres comprised smooth, highly regular particles of the expected nominal sizes (20 and 45 µm; Fig. S1A–D). By contrast, PS fragments, PS + Irgafos 168 fragments, and limestone fragments all exhibited irregular, angular shapes and heterogeneous size distributions (Fig. S1 E–J).

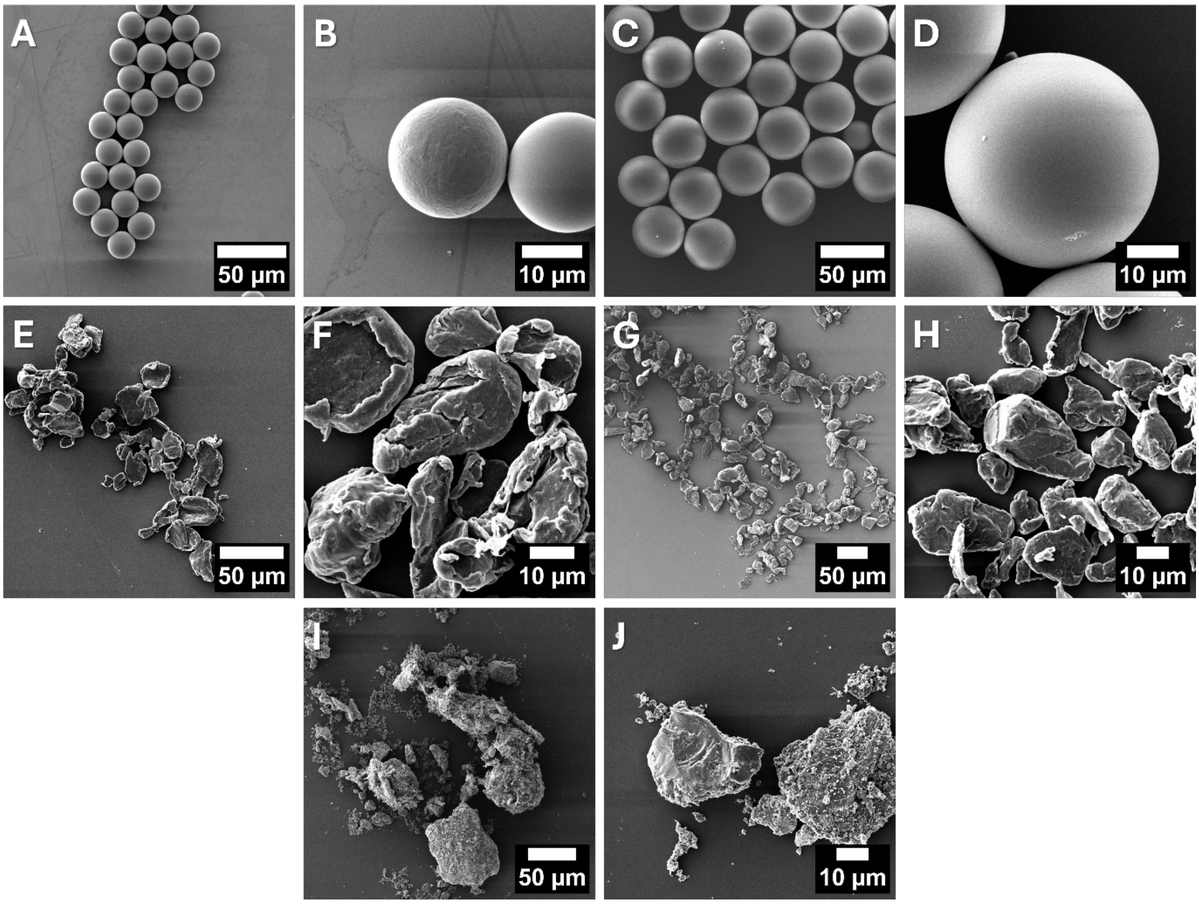


**Figure S1** Scanning electron micrographs of the used particles. 20 µm PS spheres (A, B), 45 µm PS spheres (C, D), PS fragments (E, F), PS fragments + Irgafos 168 (G, H), and Limestone fragments (I, J).
